# *Rock2* heterozygosity improves cognitive behavior and endothelial function in a mouse model of 16p11.2 deletion autism syndrome

**DOI:** 10.1101/2024.06.24.600440

**Authors:** Julie Ouellette, Baptiste Lacoste

## Abstract

Rho-associated coiled-coil containing protein kinase-2 (ROCK2) is a critical player in many cellular processes and has been incriminated in cardiovascular and neurological conditions. Recent evidence has shown that non-selective pharmacological ROCK inhibition ameliorates behavioral alterations in an autism mouse model of 16p11.2 haploinsufficiency. We had also revealed that 16p11.2-deficient mice display cerebrovascular abnormalities, including endothelial dysfunction. To investigate whether genetic blockage of ROCK2 also exerts beneficial effects on cognition and angiogenesis, we generated mice with both 16p11.2 and ROCK2 haploinsufficiency (*16p11.2*^*df/+*^*;Rock2*^*+/-*^). We find that *Rock2* heterozygosity on a *16p11.2*^*df/+*^ background rescues recognition memory and slightly reduces repetitive behaviors. Furthermore, brain endothelial cells (ECs) isolated from *16p11.2*^*df/+*^*;Rock2*^*+/-*^ mice display improved angiogenic capacity compared to ECs from *16p11.2*^*df/+*^ littermates. Overall, this study implicates *Rock2* gene as a critical modulator in 16p11.2-associated alterations, highlighting its potential as a target for treatment of autism spectrum disorders.

## INTRODUCTION

Rho-associated coiled-coil containing protein kinases (ROCK) are downstream effectors of small GTPase RhoA (1). The RhoA/ROCK pathway is involved in regulating cell migration, growth, morphogenesis and apoptosis, regulates changes in the cytoskeleton (2), and mediates vascular function (3, 4). ROCK consist of two homologous isoforms (ROCK1 and ROCK2) with ROCK2 being primarily expressed in the brain and the vasculature (1). Elevated ROCK activity is often linked with adverse effects, and reduced ROCK activity is considered beneficial in pathological conditions (5-9). For instance, increased ROCK2 activity and levels in vascular smooth muscle cells contribute to the pathogenesis of pulmonary arterial hypertension (9), while pharmacological inhibition of ROCK2 promotes vascular endothelial barrier stability in the context of respiratory distress syndrome (6). As such, a growing body of literature highlights the therapeutic potential of pharmacological ROCK inhibition (7, 8).

Considering that ROCK2 is predominant in the brain, mounting evidence demonstrates its role in neurodegeneration and more recently in neurodevelopmental disorders (7, 10). In a mouse model of 16p11.2 deletion autism syndrome, Lorenzo and colleagues demonstrated that treatment with Fasudil, a non-selective pharmacological ROCK inhibitor, improved cognitive performance in recognition and location memory (10). The 16p11.2 deletion is a common genetic risk factor for autism spectrum disorders (ASD), as this locus comprises several genes that are linked to neuropsychiatric conditions, including *Kctd13* (10, 11). *Kctd13* is crucial for late stages of brain development as it forms a complex with *Cul3* and regulates levels of RhoA that in turn control actin cytoskeletal architecture and cell motility (12). Interestingly, Lorenzo et al. also demonstrated that *Kctd13*^*+/-*^ mice phenocopied cognitive deficits identified in their mouse model of 16p11.2 deletion syndrome, and these cognitive changes were rescued by Fasudil (10). Using another mouse model of the 16p11.2 deletion syndrome, namely *16p11.2*^*df/+*^ mice (13), we have previously established a causal link between 16p11.2 deletion and altered cerebrovascular maturation, leading to lasting brain endothelial dysfunction and causing behavioral changes (14). Elevated ROCK2 activity is associated with endothelial dysfunction as it inhibits endothelial nitric oxide synthase (eNOS), which is essential for modulation of vascular tone, increases endothelial cell apoptosis, and promotes vascular hyperpermeability at the BBB (4, 6, 15). However, whether selective reduction of *Rock2* gene expression exerts beneficial effects on 16p11.2 deletion-related deficits remained to be tested. In this study, we investigate the impact of *Rock2* heterozygosity (*Rock2*^*+/-*^) on two behaviors and endothelial function in *16p11.2*^*df/+*^ mice.

## METHODS

### Animals

All animal procedures were approved by the University of Ottawa Animal Care Committee and conducted in accordance with guidelines of the Canadian Council on Animal Care

### Mouse husbandry

All mice were bred in house and housed at a maximum of five per cage with free access to water and food. All experimental animals used in this study were of a mixed B6/12/CD1 genetic background. Males *16p11.2*^*df/+*^ (Jackson laboratory, stock #013128; mixed B6/129 background) were first crossed with WT females of the same background (Jackson laboratory, stock #101043) to maintain a colony of heterozygous *16p11.2*^*df/+*^ and WT littermates. As recommended by Jackson Laboratory to improve *16p11.2*^*df/+*^ pup survival, breeding cages were supplemented with breeding chow (#2019, Envigo Teklad) and DietGel (#76A, ClearH_2_O) up to weaning age (P21). Female *Rock2*^*+/*−^ were crossed with CD1 males to maintain a separate heterozygous *Rock2*^*+/*−^ mouse colony (homozygous *Rock2* deletion is embryonically lethal). *Rock2*^*+/*−^ mice were on a standard chow diet (#2018, Envigo Teklad). *Rock2*^*+/*−^ breeders, generated on a CD1 background (16, 17), were obtained as a gift from Dr. Zhengping Jia (University of Toronto, Canada). To obtain experimental animals (*16p11.2*^*df/+*^*;Rock2*^*+/-*^), *16p11.2*^*df/+*^ males were crossed with *Rock2*^*+/*−^ females and maintained on the same diet as described for *16p11.2*^*df/+*^ mice. All assays were performed on offspring past the 5^th^ generation.

### Genotyping

*16p11.2*^*df/+*^;*Rock2*^*+/-*^ and their *16p11.2*^*df/+*^ and WT littermates were genotyped with the following primers: 5′-CCTCATGGACTAATTATGGAC-3′ (forward) and 5′-CCAGTTTCACTAATGACACA-3′ (reverse), with a PCR product of 2.2 kb for *16p11.2*^*df/+*^ mice(13). *Rock2*^*+/-*^ mice were genotyped with the following primers: 5′-CATACATGTGCCAAAATCTGCTAAC-3′ and 5′-GGGGGAACTTCCTGACTAGG-3′.

### Behavioral assays

Two-month-old adult male mice were used for this part of the study. All animals were left to acclimatize to an inverted light cycle room for 10 days prior to behavior testing. During this time, animals were handled once a day for 7 days. Behavioral tests were completed at the University of Ottawa’s Behavior Core Facility between 9:00 and 17:00 under dim red light. Before testing was started, animals were habituated in the testing room for 60 minutes. Tests were adapted from our previous work and the literature (14, 18, 19), as summarized below.

#### Novel-object recognition test

A two-day novel object recognition test was performed. On day one, each animal was habituated for 30 minutes in an empty open-field arena (45 × 45x 45cm). On the following day, experimental day two, mice were habituated in the same empty open field as in day one for 10 minutes. Once habituation was complete, each mouse was removed from the open field and placed in a clean home cage for 2 minutes. Two identical objects (red cup or white funnel) were placed in the arena, the mouse was returned and then left to explore for a 10-minute familiarization period. The mouse was then removed and placed in a clean holding cage for 1 hour. After this hour, the object-recognition test entailed one cleaned familiar object and one cleaned novel object (red cup or white funnel). The mouse was returned to the arena for a 5-minute recognition period. The interactions with the objects were recorded using Ethovision XT software (Noldus). Object recognition was scored as the amount of time the nose of the animal was within 2cm of the object. The discrimination index was calculated for the novel object as follows: [time spent interacting with novel object/ (time spent interacting with novel object + time spent interacting with familiar object)]. While the discrimination index for the familiar object was calculated as follows: [time spent interacting with familiar object/ (time spent interacting with novel object + time spent interacting with familiar object)].

#### Marble-burying test

This test entailed a 30-minute trial per mouse. Each trial was set up with a standard polycarbonate rat cage (26 × 48 x 20cm) filled with 5cm in thickness of SANI-chip bedding. A total of 20 marbles were evenly distributed (5 rows of 4 marbles) on the bedding. At the beginning of the trial, each mouse is placed in the bottom left corner and left to navigate the cage while the cage is covered by a transparent Plexiglas. After the trial, the number of marbles that were either fully buried or at least two-thirds covered by bedding were counted as buried.

### Primary mouse brain endothelial cell isolation

Cell isolation was performed as previously described (20). All mice were euthanized by cervical dislocation. The cerebral cortex was dissected in cold HBSS without calcium and magnesium using autoclaved tools which were submerged in 100% ethanol for at least 30 minutes prior to dissections. The cortex was cut in 2-3 mm pieces and dissociated in Neural Tissue Dissociation Kit P compounds (Miltenyi Biotec, 130-092-628) to obtain a cell suspension. The cell suspension was incubated with CD31-coated magnetic microbeads (Miltenyi Biotec, 130-097-418) and placed on a magnetic MACs separator to isolate endothelial cells (ECs). ECs isolated from each mouse were seeded in 2 wells of 6-well plate coated with attachment factor protein 1X (ThermoFisher Scientific, S006100). For a pure EC population, ECs were cultured in an EC specific medium (Lonza, CC-3202) which was replaced 48 hours post-seeding and every 48 hours until cells were at least 90% confluent.

### In vitro network-formation assay

EC network-formation assays were performed using growth-factor-reduced Matrigel (BD Bioscience, cat. no. C354230). Growth-factor-reduced condition includes: EGF < 0.5 ng ml–1, PDGF < 5 pg ml–1, IGF1 = 5 ng ml–1 and TGF-β = 1.7 ng ml–1. Each well of a 96-well plate was coated with 50 μl of Matrigel and incubated at 37 °C for 30 minutes to promote polymerization. ECs were collected with TryplE (Gibco, 12604013) and counted using a hemocytometer. A total of 2 × 10^4^ cells were seeded in each well with 150 μl EC specific media (Lonza, CC-3202). TIFF images of capillary-like networks were captured using a Zeiss Axio Image M2 microscope equipped with a digital camera at 4, 8, and 24h post-seeding. Images were processed using the Angiogenesis function of ImageJ (21).

### Statistical analyses

Sample sizes are similar to previously published report (14). A power analysis was not performed to determine sample sizes. All replication experiments were successful. All data analysis was conducted blind to genotype and groups were reassembled following data analysis. All animals were numbered to maintain randomization. Statistical tests were performed using GraphPad Prism 10.0 Software. Data distribution was assumed to be normal, but this was not formally tested. A two-way analysis of variance (ANOVA; e.g. ‘genotype x object’ or ‘genotype x time’), and a *post-hoc* test (Tukey’s multiple comparisons) was used for the novel object recognition test and *in vitro* network formation assay. A one-way analysis of variance (ANOVA), and a *post-hoc* test (Tukey’s multiple comparisons) was used for marble burying test. *P*<0.05 was considered significant. Statistical details can be found in the figure legend.

## RESULTS

To determine whether *Rock2* heterozygosity improves 16p11.2 deletion-associated cognitive changes, we subjected *16p11.2*^*df/+*^;*Rock2*^*+/-*^ mice and their control littermates (WT, *16p11.2*^*df/+*^) to two behavioral tests: a novel object recognition task, or NOR (assessing recognition memory), and a marble burying task (assessing repetitive behaviors). In the NOR, consistent with our previous findings (14), *16p11.2*^*df/+*^ mice exhibited a significant preference for the familiar object as opposed to the novel object. However, compound *16p11.2*^*df/+*^;*Rock2*^*+/-*^ mice displayed a similar discrimination index to that of their WT littermates. Notably, *16p11.2*^*df/+*^;*Rock2*^*+/-*^ mice had an increased discrimination index for the novel object compared to *16p11.2*^*df/+*^ littermates (Fig. 1A). While, as expected, *16p11.2*^*df/+*^ mice buried significantly more marbles, *16p11.2*^*df/+*^;*Rock2*^*+/-*^ mice showed a reduction, albeit non-significant, in repetitive behaviors, at levels similar to WT controls (Fig. 1B). These results suggest that *Rock2* heterozygosity has the potential to improve 16p11.2-deletion induced behavioral phenotypes.

**Fig 1.**
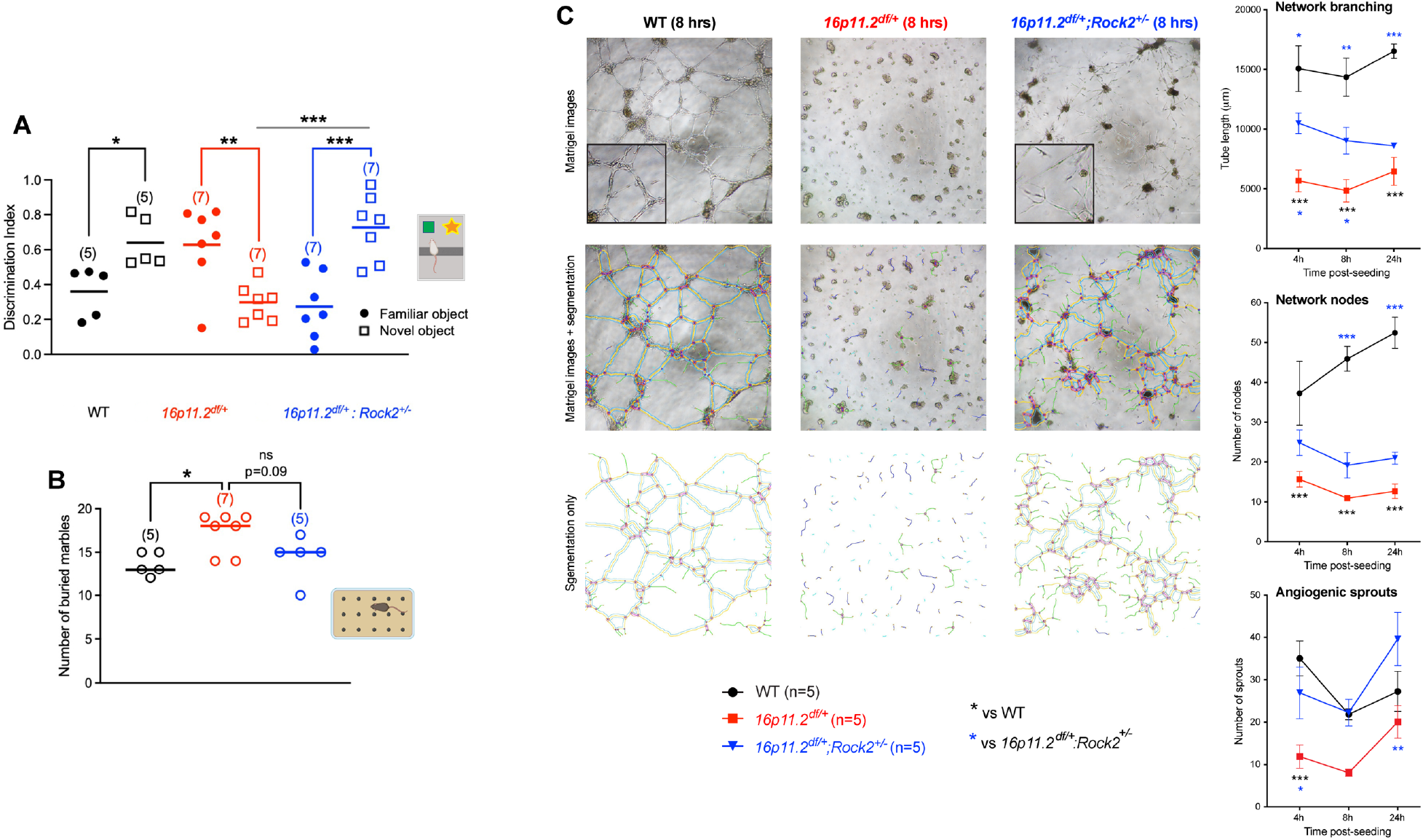
*Rock2* heterozygosity improves 16p11.2 deletion-associated behavioral deficits and brain endothelial cell dysfunction. *A,B*, Behavioral assessment of *16p11.2*^*df/+*^*;Rock2*^*+/-*^, *16p11.2*^*df/+*^ and WT littermates at P50. *A*, Novel object recognition task revealed improved recognition memory in *16p11.2*^*df/+*^*;Rock2*^*+/-*^ mice compared to *16p11.2*^*df/+*^ littermates. *B*, Marble burying task demonstrated a slight decrease in the number of marbles buried by *16p11.2*^*df/+*^*;Rock2*^*+/-*^ mice compared to *16p11.2*^*df/+*^ littermates. *C, Left*, representative images of the tube formation assay 8 hours after seeding primary brain endothelial cells, with respective network traces below. We identified increased capillary-like networks in *16p11.2*^*df/+*^*;Rock2*^*+/-*^ ECs compared to ECs from *16p11.2*^*df/+*^ littermates. *Right*, quantification of network branching (*top*), network nodes (*middle*), and angiogenic sprouts (*bottom*). Data are mean ± s.e.m. ns: not significant, *P<0.05, **P<0.01, ***P<0.001 (two-way ANOVA and Tukey’s *post-hoc* test *in A,C;* one-way ANOVA and Tukey’s *post-hoc* test in *B*).

We have previously reported that primary brain endothelial cells (ECs) isolated from *16p11.2*^*df/+*^ mice are unable to form a capillary-like network *in vitro* (14). Here, we sought to assess the impact of *Rock2* heterozygosity on brain endothelial function. We seeded primary brain ECs from WT, *16p11.2*^*df/+*^ and *16p11.2*^*df/+*^*;Rock2*^*+/-*^ littermates in a growth factor reduced Matrigel^®^ and evaluated capillary-like network formation 4-, 8- and 24-hours after cell seeding. ECs from WT mice showed an expected increase in tube length and node number (Fig. 1C). In agreement with our previous study (14), 16p11.2-deficient ECs were unable to form an extensive capillary-like network when compared to WT ECs (Fig. 1C). From 4- to 24-hours post-seeding, little to no network was formed by 16p11.2-deficient ECs despite a slight increase at 24 hours (Fig. 1C). Interestingly, *16p11.2*^*df/+*^*;Rock2*^*+/-*^ ECs showed an overall increased ability to form capillary-like structures compared to 16p11.2-deficient ECs (Fig. 1C). We measured a significant increase in tube length at 4- and 8-hours post-seeding compared to ECs isolated from *16p11.2*^*df/+*^ littermates (Fig. 1C). At 24 hours post-seeding, *16p11.2*^*df/+*^*;Rock2*^*+/-*^ EC tube length returned closer to *16p11.2*^*df/+*^ ECs network values, consistent with lack of trophic support towards the end of the assay (Fig. 1C). Compared to *16p11.2*^*df/+*^ ECs, a non-significant increase in network nodes was measured in *16p11.2*^*df/+;*^*Rock2*^*+/-*^ EC throughout the 24-hour assay (Fig. 1C). At 24 hours, *16p11.2*^*df/+*^*;Rock2*^*+/-*^ EC network branching decreased while WT and *16p11.2*^*df/+*^ ECs increased network branching (Fig. 1C). For network nodes, *16p11.2*^*df/+*^*;Rock2*^*+/-*^ and *16p11.2*^*df/+*^ ECs had similar values whereas WT network nodes consistently increased throughout the 24-hour assay (Fig. 1C). Finally, ROCK2 heterozygosity in *16p11.2*^*df/+*^ ECs increased the number of angiogenic sprouts 4- and 24-hours post-seeding compared to *16p11.2*^*df/+*^ ECs, suggesting an ameliorating of angiogenic initiation (Fig. 1C). These results demonstrate that *Rock2* heterozygosity can partially alleviate 16p11.2 deletion-associated brain endothelial dysfunction.

## DISCUSSION

In this study, we provide further evidence supporting the concept that reducing ROCK2 levels/activity might represent a valid therapeutic avenue for neurodevelopmental disorders. We show that *Rock2* heterozygosity can ameliorates key features in 16p11.2-deficient mice, namely recognition memory and endothelial function.

The RhoA/ROCK pathway is a target in various central nervous system disorders as pharmacological modulation of ROCK activity is increasingly recognized as valid disease-modifying strategy (7, 10). For example, ROCK inhibitor Fasudil can modulate the aggregation of α-synuclein and ameliorate motor behavior in a mouse model of Parkinson’s disease (22). In an *in vitro* Alzheimer’s disease (AD) model, ROCK2 inhibitor TA8Amino increased neurogenesis and neuritogenesis. As a result, ROCK inhibition was proposed as a possible therapy for AD to target reduced synapses and decreased adult neurogenesis induced by accumulation of hyperphosphorylated tau protein (23). In addition, a rat model of delayed encephalopathy after acute carbon monoxide poisoning (DEACMP) displayed elevated ROCK2 protein level. Following inhibition of ROCK2 with Y-27632 for three weeks, DEACMP rats showed reduced neuronal damage, increased myelin repair and improved cognitive and motor ability (24).

Moreover, mutations in *Ccm1,-2*, or *-3* is associated with vascular endothelial cell defects often leading to cerebral cavernous malformations (CCM), lesions causing seizures and stroke. In cells lines of CCM1,-2, or -3 knockdown, RhoA was found to be overexpressed with persistent activity. CCM1,-2, or -3 knockdown cell lines also revealed an inability to form vessel-like tubes *in vitro*. Following treatment of ROCK inhibitor Y-27632 or ROCK2 knockdown, tube formation was rescued in this model (25).

Here, we provide further evidence supporting the involvement of the RhoA/ROCK pathway in neurodevelopmental disorders. Data obtained with *16p11.2*^*df/+*^*;Rock2*^*+/-*^ mice reveal that developmental reduction of *Rock2* expression triggers long-term beneficial effects in this model, such as rescued recognition memory. The significantly improved recognition memory in *16p11.2*^*df/+*^*;Rock2*^*+/-*^ mice is in line with a recent study demonstrating that chronic ROCK inhibition with Fasudil ameliorated cognitive function in 16p11.2 deletion mice (10). Beneficial effects of *Rock2* ablation may be related to decreased RhoA signaling as *Kctd13* expression, a 16p11.2 locus gene regulator of RhoA protein level, is reduced (26). Without regulation of the RhoA/ROCK pathway via *Kctd13*, elevated RhoA/ROCK activity could be pathogenic and may be involve in regulating 16p11.2 deletion-associated phenotypes.

The improved angiogenic response of *16p11.2*^*df/+*^*;Rock2*^*+/-*^ ECs highlights the beneficial properties of decreased ROCK2 in endothelial cells. In agreement with our results, a previous study using a model of hindlimb ischemia showed enhanced endothelial tube formation using a ROCK inhibitor (27). Increased endothelial branching and sprouting measured in *16p11.2*^*df/+*^*;Rock2*^*+/-*^ ECs may be in part attributed to role of the RhoA/ROCK pathway in cytoskeletal reorganization (28). While the role of ROCK in angiogenesis is not fully elucidated, it was suggested that vascular endothelial growth factor (VEGF) is involved in promoting EC migration and proliferation (29).

In our prior work, we had demonstrated that endothelium-specific 16p11.2 haploinsufficiency in mice was sufficient to drive ASD-related behaviors (14). Here, we show an improved endothelial function in 16p11.2-deficient mice with *Rock2* heterozygosity, raising the possibility that improved behaviors in *16p11.2*^*df/+*^*;Rock2*^*+/-*^ mice may result, at least in part, from improved vascular function. However, since we do not provide a direct causal link, one must consider the influence of *Rock2* heterozygosity on other factors and/or cell-types that could contribute to rescued cognitive performance and endothelial function. For instance, pharmacological inhibition of ROCK exerts a pro-survival effect on neurons and can prevents learning and memory impairments (30, 31). Moreover, in cortical organoids derived from 16p11.2 deletion carriers, ROCK inhibition rescued delayed migration of neurons (32). In a mouse model deficient for *Kctd13*, increased RhoA levels were associated with altered synaptic transmission, which was recovered following ROCK inhibition (33). Taken together, *Rock2* heterozygosity in 16p11.2-deficient mice may not only impact the vasculature but also neuronal cells, in turn improving behavior in this model. Future research is required to unmask the mechanisms underlying such rescue.

Overall, this work highlights the beneficial aspects of *Rock2* heterozygosity in an ASD mouse model, further supporting the RhoA/ROCK pathway as a potential therapeutic target for neurodevelopmental disorders.

## Acknowledgments

This research was funded by two grants from the Canadian Institutes of Health Research (#388805,# 506513) to B.L.; a Scholarship from the Canadian Institutes of Health Research to J.O.

## REFERENCES

1. Noma K, Kihara Y, Higashi Y. Striking crosstalk of ROCK signaling with endothelial function. J Cardiol. 2012;60(1):1–6.

2. Hall A. Rho family GTPases. Biochem Soc Trans. 2012;40(6):1378–82.

3. Strassheim D, Gerasimovskaya E, Irwin D, Dempsey EC, Stenmark K, Karoor V. RhoGTPase in Vascular Disease. Cells. 2019;8(6).

4. Yao L, Romero MJ, Toque HA, Yang G, Caldwell RB, Caldwell RW. The role of RhoA/Rho kinase pathway in endothelial dysfunction. J Cardiovasc Dis Res. 2010;1(4):165–70.

5. Rikitake Y, Liao JK. ROCKs as therapeutic targets in cardiovascular diseases. Expert Rev Cardiovasc Ther. 2005;3(3):441–51.

6. Beckers CM, Knezevic N, Valent ET, Tauseef M, Krishnan R, Rajendran K, et al. ROCK2 primes the endothelium for vascular hyperpermeability responses by raising baseline junctional tension. Vascul Pharmacol. 2015;70:45–54.

7. Koch JC, Tatenhorst L, Roser AE, Saal KA, Tonges L, Lingor P. ROCK inhibition in models of neurodegeneration and its potential for clinical translation. Pharmacol Ther. 2018;189:1–21.

8. Weber AJ, Herskowitz JH. Perspectives on ROCK2 as a Therapeutic Target for Alzheimer’s Disease. Front Cell Neurosci. 2021;15:636017.

9. Shimizu T, Fukumoto Y, Tanaka S, Satoh K, Ikeda S, Shimokawa H. Crucial role of ROCK2 in vascular smooth muscle cells for hypoxia-induced pulmonary hypertension in mice. Arterioscler Thromb Vasc Biol. 2013;33(12):2780–91.

10. Martin Lorenzo S, Nalesso V, Chevalier C, Birling MC, Herault Y. Targeting the RHOA pathway improves learning and memory in adult Kctd13 and 16p11.2 deletion mouse models. Mol Autism. 2021;12(1):1.

11. Crepel A, Steyaert J, De la Marche W, De Wolf V, Fryns JP, Noens I, et al. Narrowing the critical deletion region for autism spectrum disorders on 16p11.2. Am J Med Genet B Neuropsychiatr Genet. 2011;156(2):243–5.

12. Lin P, Yang J, Wu S, Ye T, Zhuang W, Wang W, et al. Current trends of high-risk gene Cul3 in neurodevelopmental disorders. Frontiers in Psychiatry. 2023;14.

13. Horev G, Ellegood J, Lerch JP, Son YE, Muthuswamy L, Vogel H, et al. Dosage-dependent phenotypes in models of 16p11.2 lesions found in autism. Proc Natl Acad Sci U S A. 2011;108(41):17076–81.

14. Ouellette J, Toussay X, Comin CH, Costa LDF, Ho M, Lacalle-Aurioles M, et al. Vascular contributions to 16p11.2 deletion autism syndrome modeled in mice. Nat Neurosci. 2020;23(9):1090–101.

15. Huang L, Dai F, Tang L, Bao X, Liu Z, Huang C, et al. Distinct Roles For ROCK1 and ROCK2 in the Regulation of Oxldl-Mediated Endothelial Dysfunction. Cell Physiol Biochem. 2018;49(2):565–77.

16. Soliman H, Nyamandi V, Garcia-Patino M, Varela JN, Bankar G, Lin G, et al. Partial deletion of ROCK2 protects mice from high-fat diet-induced cardiac insulin resistance and contractile dysfunction. Am J Physiol Heart Circ Physiol. 2015;309(1):H70–81.

17. Zhou Z, Meng Y, Asrar S, Todorovski Z, Jia Z. A critical role of Rho-kinase ROCK2 in the regulation of spine and synaptic function. Neuropharmacology. 2009;56(1):81–9.

18. Angoa-Pérez M, Kane MJ, Briggs DI, Francescutti DM, Kuhn DM. Marble Burying and Nestlet Shredding as Tests of Repetitive, Compulsive-like Behaviors in Mice. J Vis Exp. 2013(82):e50978.

19. Portmann T, Yang M, Mao R, Panagiotakos G, Ellegood J, Dolen G, et al. Behavioral abnormalities and circuit defects in the basal ganglia of a mouse model of 16p11.2 deletion syndrome. Cell Rep. 2014;7(4):1077–92.

20. Ouellette J, Lacoste B. Isolation and functional characterization of primary endothelial cells from mouse cerebral cortex. STAR Protoc. 2021;2(4):101019.

21. Carpentier G. Angiogenesis Analyzer for ImageJ 2012 [Available from: http://image.bio.methods.free.fr/ImageJ/?Angiogenesis-Analyzer-for-ImageJ#outil_sommaire_0.

22. Tatenhorst L, Eckermann K, Dambeck V, Fonseca-Ornelas L, Walle H, Lopes da Fonseca T, et al. Fasudil attenuates aggregation of alpha-synuclein in models of Parkinson’s disease. Acta Neuropathol Commun. 2016;4:39.

23. Moreira N, Tamarozzi ER, Lima J, Piassi LO, Carvalho I, Passos GA, et al. Novel Dual AChE and ROCK2 Inhibitor Induces Neurogenesis via PTEN/AKT Pathway in Alzheimer’s Disease Model. Int J Mol Sci. 2022;23(23).

24. Xu L, Liu X, Guo C, Wang C, Zhao J, Zhang X, et al. Inhibition of ROCK2 kinase activity improved behavioral deficits and reduced neuron damage in a DEACMP rat model. Brain Res Bull. 2022;180:24–30.

25. Borikova AL, Dibble CF, Sciaky N, Welch CM, Abell AN, Bencharit S, et al. Rho kinase inhibition rescues the endothelial cell cerebral cavernous malformation phenotype. J Biol Chem. 2010;285(16):11760–4.

26. Richter M, Murtaza N, Scharrenberg R, White SH, Johanns O, Walker S, et al. Altered TAOK2 activity causes autism-related neurodevelopmental and cognitive abnormalities through RhoA signaling. Mol Psychiatry. 2018.

27. Fayed HS, Bakleh MZ, Ashraf JV, Howarth A, Ebner D, Al Haj Zen A. Selective ROCK Inhibitor Enhances Blood Flow Recovery after Hindlimb Ischemia. Int J Mol Sci. 2023;24(19).

28. Amano M, Nakayama, M. and Kaibuchi, K. Rho-Kinase/ROCK: A Key Regulator of the Cytoskeleton and Cell Polarity. Cytoskeleton. 2010;67:545–56.

29. Bryan BA, Dennstedt E, Mitchell DC, Walshe TE, Noma K, Loureiro R, et al. RhoA/ROCK signaling is essential for multiple aspects of VEGF-mediated angiogenesis. FASEB J. 2010;24(9):3186–95.

30. Tonges L, Koch JC, Bahr M, Lingor P. ROCKing Regeneration: Rho Kinase Inhibition as Molecular Target for Neurorestoration. Front Mol Neurosci. 2011;4:39.

31. Bobo-Jimenez V, Delgado-Esteban M, Angibaud J, Sanchez-Moran I, de la Fuente A, Yajeya J, et al. APC/C(Cdh1)-Rock2 pathway controls dendritic integrity and memory. Proc Natl Acad Sci U S A. 2017;114(17):4513–8.

32. Urresti J, Zhang P, Moran-Losada P, Yu NK, Negraes PD, Trujillo CA, et al. Cortical organoids model early brain development disrupted by 16p11.2 copy number variants in autism. Mol Psychiatry. 2021;26(12):7560–80.

33. Sundberg M, Pinson H, Smith RS, Winden KD, Venugopal P, Tai DJC, et al. 16p11.2 deletion is associated with hyperactivation of human iPSC-derived dopaminergic neuron networks and is rescued by RHOA inhibition in vitro. Nat Commun. 2021;12(1):2897.

